# Systemic delivery of CRISPR-Cas9 nickase suppresses oncogene amplified cancer progression

**DOI:** 10.64898/2026.04.26.720919

**Authors:** Matthew B. Hanlon, Scot A. Wolfe

**Affiliations:** Department of Molecular, Cell and Cancer Biology, University of Massachusetts Chan Medical School, Worcester, MA 01605, USA; Li Weibo Institute for Rare Diseases Research, University of Massachusetts Chan Medical School, Worcester, MA, USA

## Abstract

Oncogene amplification is a key driver of tumorigenesis and a perpetuator of genomic instability. Oncogene amplification accelerates cancer cell proliferation and evolution, contributing substantially to the enhancement of adaptation mechanisms, such as treatment resistance, which pose a significant therapeutic challenge. However, previous studies have shown oncogene amplification to be a critical vulnerability, rendering cancer cells, but not normal cells, susceptible to targeted, CRISPR-Cas9 nickase – mediated DNA damage and cell death *in vitro*. Here, we demonstrate the initial framework for the translation of this potential therapeutic approach utilizing Cas9^D10A^ – mRNA and functionalized lipid nanoparticles for the targeted delivery, and suppression of disseminated *MYCN*-amplified neuroblastoma *in vivo*.

---

Oncogene amplification is a common genomic phenomenon observed in a multitude of cancers and is typically associated with advanced stage disease ^1–4^. Gene amplification is a copy number increase of a chromosomal region harboring one or more genes and regulatory elements ^3,5,6^. Often spanning from hundreds of kilobases (kb) to several megabases (Mb) ^3,7–9^, amplified genomic regions can be organized into contiguous units within the linear genome, commonly referred to as a homogenously staining region (HSR), or as multi-copy, circular extrachromosomal DNA (ecDNA) structures ^1,10–12^. Inherent to the cancer genome, gene amplification is reasoned to be a consequence of genomic instability. Genomic instability is an increased tendency to acquire both small and large-scale genomic alterations through defects in DNA repair machinery, replication licensing, or cell-cycle control ^13–15^. Working in tandem, oncogene amplification and genomic instability accelerate clonal evolution through enhanced proliferation and genomic heterogeneity ^16^. Consequentially, the enhanced adaptive potential of oncogene-amplified cancers poses a substantial therapeutic challenge, as conventional treatments impose selective pressures that promote the development of resistance mechanisms ^13,15–19^. Neuroblastoma, an aggressive pediatric cancer of the sympathetic nervous system, exemplifies a difficult to treat, oncogene-amplified cancer ^20,21^. Currently, neuroblastoma accounts for ∼15% of all childhood cancer mortality ^21–23^.

Neuroblastomas frequently present with a high degree of genomic heterogeneity, such as amplification of the *MYCN* oncogene, which is observed in ∼25% of cases ^20,24,25^. *MYCN* amplifications are highly variable, demonstrating copy number gains of tens to thousands where each amplified segment spans ∼350 kb – 8 Mb in length ^9,26–28^. Importantly, *MYCN*-amplification is strongly correlated with high-risk disease and a poor prognosis ^24^. Currently, the standard of care for high-risk neuroblastoma (HR-NB) is an intensive, multimodal regimen that incorporates high-dose chemotherapy, radiation, myeloablation with hematopoietic stem cell transplant, and immunotherapy ^20,23^. Despite aggressive treatment, overall survival for HR-NB is ∼50% with a high probability of relapse, 80% of which occur within 2-years of diagnosis ^20,23,29,30^. Currently, there is no standard of care or salvage therapy known to be curative for relapsed or refractory neuroblastoma ^20,29,31^, highlighting the urgent need for novel therapeutic interventions.

Previously, we demonstrated that CRISPR-Cas9 nickases can selectively promote cell death in a gene amplification-dependent manner *in vitro* ^32^. Cas9 nickase, which has one of its two DNA cleavage domains (HNH or RuvC) inactivated, generates a targeted single-strand break (SSB) in DNA ^33–35^. Single-strand breaks can occur within the genome of a cell through a variety of processes and exhibit low intrinsic toxicity, particularly in post-mitotic cells where DNA replication is not actively occurring ^36^. However, in proliferating cells, the accumulation of SSBs can inflict substantial cellular toxicity ^32,36–38^. If left unrepaired, SSBs can be converted into toxic single-ended double-strand breaks (seDSBs) upon collision with a DNA replication fork inducing fork collapse, replication stress, and cell death when present in large numbers ^32,36,39,40^.

Using *MYCN*-amplified neuroblastoma as a model system, we demonstrate the utility of CRISPR-Cas9 nickases as a potential genotype-specific therapeutic for oncogene-amplified cancers *in vivo*. Oncogene amplification is frequently associated with metastatic progression ^4,41–43^ requiring systemic interventions ^44^. Lipid nanoparticles (LNPs) have emerged as a promising platform for the targeted delivery of novel RNA-based therapeutics via systemic administration ^45–49^. Taking full advantage of this delivery platform’s modularity, we employ functionalized LNP carriers for the systemic, targeted delivery of mRNA payloads to primary *MYCN*-amplified neuroblastoma tumors *in vivo*. Importantly, we demonstrate potent activity of LNP-delivered Cas9 nickase in a disseminated tumor model with promising therapeutic efficacy.

## RESULTS

### Assessing the passive biodistribution of candidate LNP formulations

For delivery of nucleic acids, LNPs consist of four essential components; 1) ionizable or cationic lipids for the efficient packaging of cargo; 2) helper phospholipids and 3) cholesterol for structural stability and endosomal escape; and 4) PEGylated lipids for colloidal stability, improved circulation time, and reduced serum protein interaction ^50–52^. Referred to as passive targeting, different lipid compositions confer unique physical properties that influence uptake in different tissues ^53,54^. As such, we evaluated the passive biodistribution of two ionizable cationic (DLin-MC3-DMA ^55^ & 306-O12B ^56^) and one permanently cationic (DOTAP ^57^) LNP formulations by assessing their uptake in primary neuroblastoma tumors (Fig. 1). The four-component ionizable cationic LNP formulations evaluated comprise DLin-MC3-DMA, DSPC, cholesterol and DMG-PEG_2000_; or 306-O12B, DOPC, cholesterol and DMG-PEG_2000_ (Fig. 1A). The four-component permanently cationic LNP formulation evaluated comprise DOTAP, DSPC, cholesterol and DMG-PEG_2000_ (Fig. 1A). All LNP formulations integrated these components at a molar ratio of 50/10/38.5/1.5, respectively.

**Figure 1.**
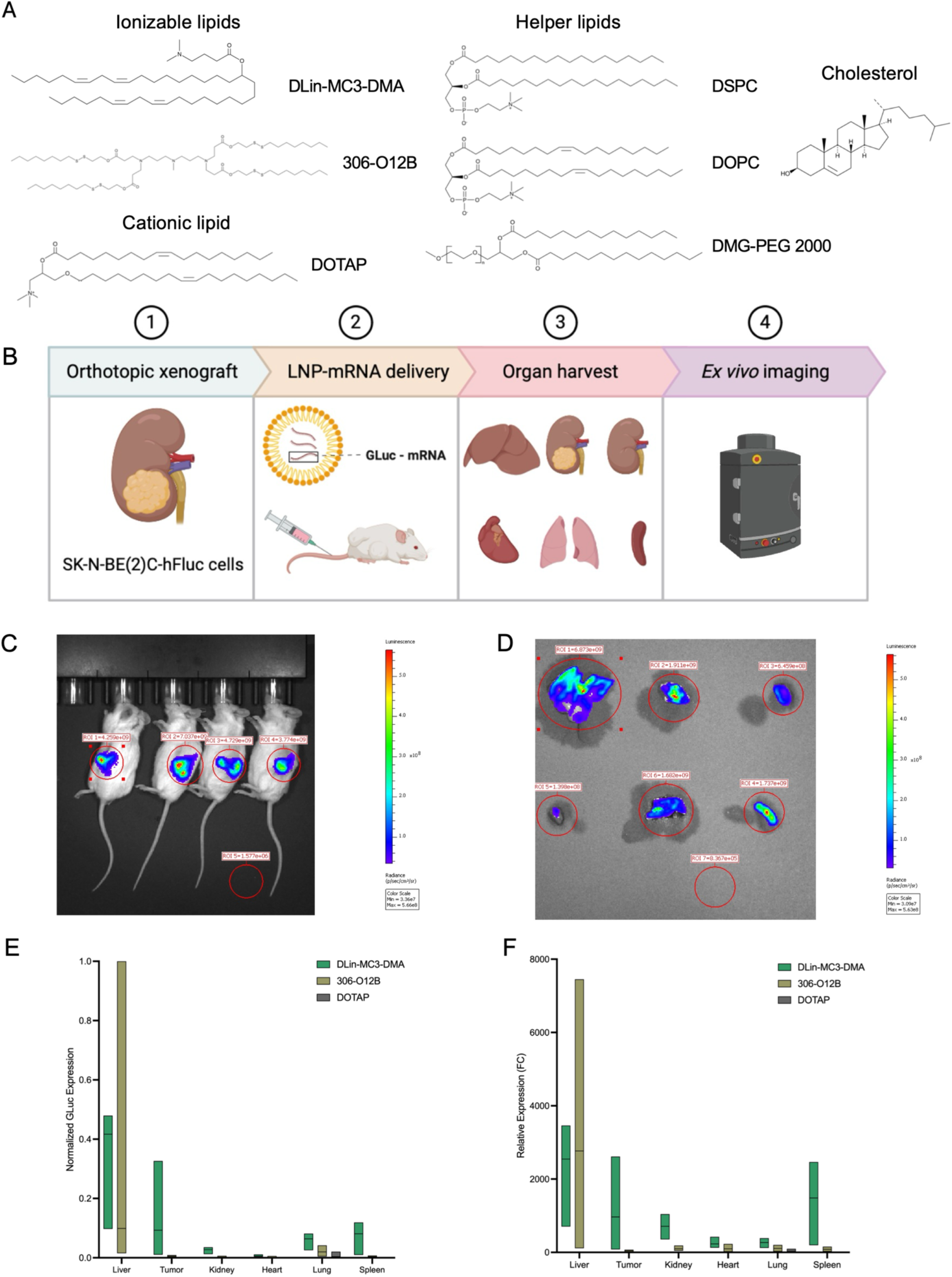
DLin-MC3-DMA LNPs demonstrate superior tumor uptake compared to 306-O12B and DOTAP LNPs. **A)** Chemical structures of lipid components used to generate LNPs for the delivery of mRNA, including ionizable cationic lipids DLin-MC3-DMA and 306-O12B, permanently cationic lipid DOTAP, helper phospholipids DSPC and DOPC, cholesterol, and DMG-PEG_2000_. Images were adapted from Cayman Chemical for noncommercial reference. **B)** Experimental design to assess the passive targeting of LNP formulations to the orthotopically engrafted SK-N-BE(2)C tumor and other organs. **C)** Representative *in vivo* bioluminescence image of orthotopically engrafted mice 12-days post-engraftment. **D)** Representative *ex vivo* bioluminescence image of mouse organs 24-hours post-delivery of LNPs encapsulating GLuc mRNA. Top row (left to right): liver, tumor, contralateral kidney. Bottom row (left to right): heart, lungs, spleen. **E – F)** Normalized BLI of each organ treatment group to the BLI of all organs combined (E) and organ-specific BLI of each treatment group relative to a vehicle-only organ control (F) as determined by *ex vivo* bioluminescence imaging of mouse organs at 24-hours post-delivery of DLin-MC3-DMA (n = 5 mice), 306-O12B (n = 3 mice), and DOTAP (n = 6 mice) LNPs encapsulating GLuc mRNA (3 mg/kg), where each group is baseline-corrected to a vehicle only control (n = 6 mice) and the mean of the population is indicated as a line.

For the initial assessment of LNP biodistribution, we orthotopically implanted human, *MYCN*-amplified neuroblastoma cells (SK-N-BE(2)C), which were modified to express firefly luciferase-P2A-mCherry as a stable transgene for disease monitoring, into the renal capsule of immunocompromised NOD SCID gamma (NSG) mice (Fig. 1B). Primary tumors were allowed to develop over 10 – 12 days and were confirmed by *in vivo* bioluminescence imaging (Fig. 1C). Next, we encapsulated *Gaussia* luciferase (GLuc) mRNA in DLin-MC3-DMA, 306-O12B, and DOTAP LNPs and characterized their uniformity by dynamic light scattering (DLS). Resulting LNPs demonstrated average diameters (Z-avg) of 88, 75, and 85 nm, respectively, and polydispersity indexes (PDI) < 0.2. We administered these GLuc mRNA LNPs to mice intravenously (IV; 3 mg/kg mRNA; Fig. 1B) and organs were harvested 24-hours post-injection. Here, the biodistribution of the LNP carriers was ascertained by quantifying GLuc expression, by means of bioluminescence intensity (BLI), in the liver, primary tumor, contralateral kidney, heart, lungs, and spleen by *ex vivo* bioluminescence imaging (Fig. 1B & D).

Both 306-O12B and DOTAP LNPs displayed negligible uptake in the primary tumor, rather a narrow biodistribution with localized expression in the liver and lungs, respectively (Fig. 1 D, E & F). However, DLin-MC3-DMA LNPs displayed a much broader biodistribution with a high degree of liver uptake, as anticipated, as well appreciable uptake in the primary tumor and adjacent tissues (Fig. 1 D, E & F). Informed by these preliminary results, we selected DLin-MC3-DMA LNPs to serve as the foundational carrier system for the initial development of a Cas9 nickase – mRNA delivery platform for neuroblastoma.

### Functionalization of LNPs for active targeting and enhanced uptake in neuroblastoma

Given the appreciable biodistribution and uptake of DLin-MC3-DMA LNPs at baseline, we next investigated surface modifications as a means of active targeting ^54^ for enhanced uptake in neuroblastoma tumors. Disialoganglioside GD2 is a surface glycolipid that is significantly overexpressed in neuroblastomas ^58^ while only being expressed at very low levels in neurons, melanocytes, and peripheral sensory nerve fibers ^59^. As such, GD2 has been classified as a tumor-associated antigen (TAA) and is a prime target of immunotherapies used in the treatment of neuroblastoma ^60–62^. One prominent example being ch14.18 (Dinutuximab), a monoclonal antibody (mAb) commonly used to treat HR-NB ^60,63^.

We reasoned ch14.18 could be utilized as a targeting moiety to enhance neuroblastoma-specific uptake of LNPs and reduce non-specific interactions with other tissues. First, we modified the DLin-MC3-DMA formulation to incorporate a maleimide functionalized PEG with a PEG:PEG-maleimide ratio of 5:1. Next, we modified ch14.18 with SATA (N-succinimidyl S-acetylthioacetate), introducing sulfhydryl groups for thiol-maleimide conjugation (Fig. 2A) ^64^. Successful conjugation was determined by size exclusion chromatography (SEC) and dynamic light scattering (DLS) demonstrating an increase in LNP diameter of ≥20 nm following mAb conjugation (Fig. 2B, Sup. Fig. 1A & B), which is consistent with previous descriptions ^65^.

**Figure 2.**
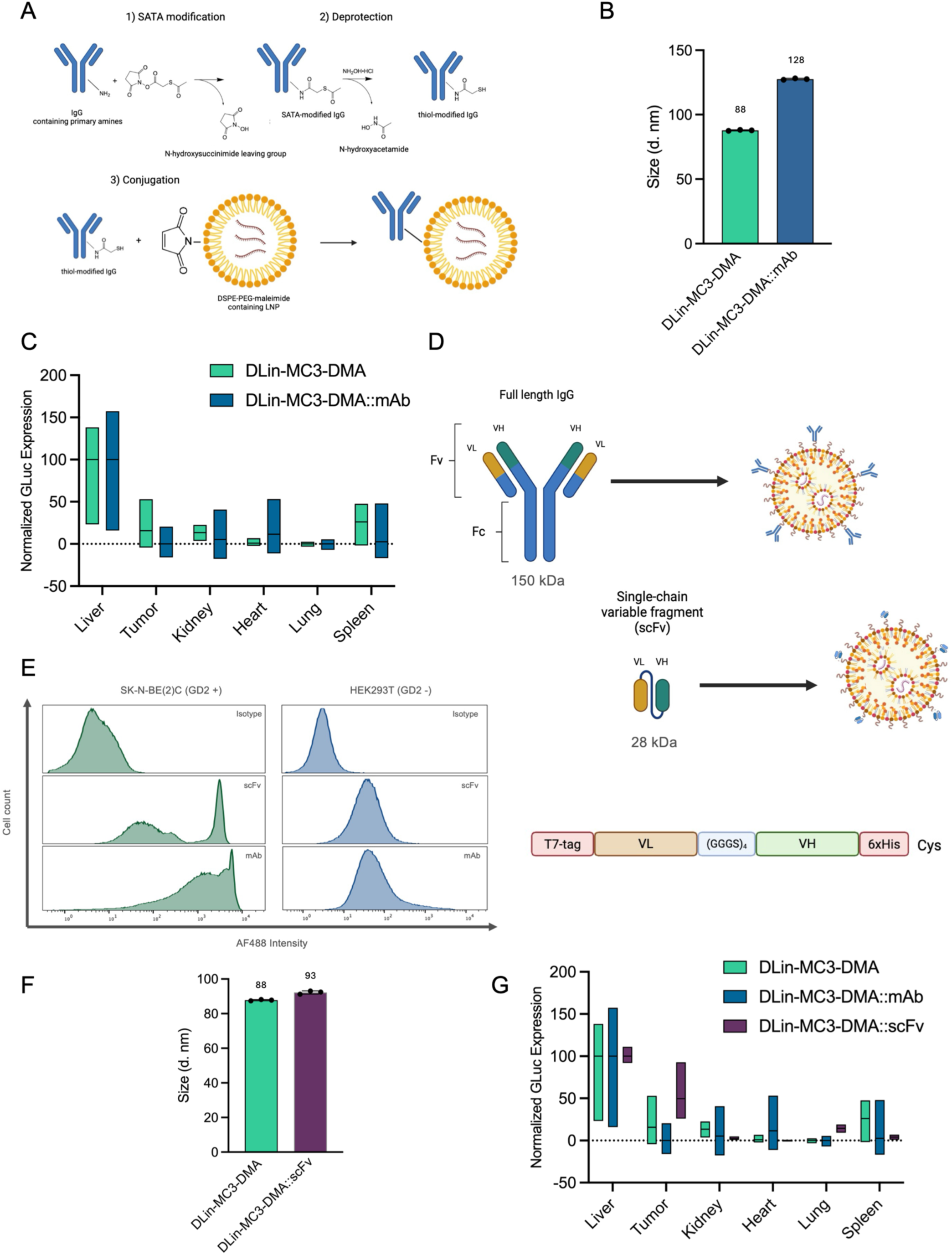
Functionalization of DLin-MC3-DMA LNPs with anti-GD2 mAb or scFv to improve neuroblastoma uptake. **A)** Schematic depicting the SATA modification of anti-GD2 antibodies with subsequent thiol-maleimide conjugation to generate antibody-coated LNPs. **B)** Average size (Z-avg) of DLin-MC3-DMA and DLin-MC3-DMA – antibody conjugated (DLin-MC3-DMA::mAb) LNPs encapsulating GLuc mRNA (3 mg/kg) as determined by dynamic light scattering (DLS). Data are presented as mean ± s.d. **C)** Normalized GLuc expression as determined by *ex vivo* bioluminescence imaging of mouse organs at 24-hours post-delivery of DLin-MC3-DMA (n = 5 mice) and DLin-MC3-DMA::mAb (n = 5 mice) LNPs containing GLuc mRNA (3 mg/kg) and baseline-corrected to a vehicle only control (n = 7 mice). Data are presented with a line indicating the mean. **D)** Diagram demonstrating the difference in size and structure of IgG (top) and scFv (middle), and a schematic corresponding to the composition of the engineered scFv (bottom). **E)** Representative histogram corresponding to the flow cytometric analysis of anti-GD2 mAb and anti-GD2 scFv activity in GD2 positive, SK-N-BE(2)C, and GD2 negative, HEK293T cells (n = 3 independent replicates). **F)** Average size (Z-avg) of DLin-MC3-DMA and DLin-MC3-DMA – scFv conjugated (DLin-MC3-DMA::scFv) LNPs encapsulating GLuc mRNA (3 mg/kg) as determined by DLS. Data is presented as mean ± s.d. **G)** Normalized GLuc expression as determined by *ex vivo* bioluminescence imaging of mouse organs at 24-hours post-delivery of DLin-MC3-DMA (n = 5 mice), DLin-MC3-DMA::mAb (n = 5 mice), and DLin-MC3-DMA::scFv (n = 3 mice) LNPs containing GLuc mRNA (3 mg/kg) and baseline-corrected to a vehicle only control (n = 5 mice). Data are presented with a line indicating the mean.

Using the same orthotopic neuroblastoma mouse model, LNP-GD2 mAb conjugates carrying GLuc mRNA (3 mg/kg mRNA) were administered intravenously. Organs were harvested 24-hours post-injection to assess GLuc expression, by means of BLI, in the liver, primary tumor, contralateral kidney, heart, lungs, and spleen by *ex vivo* bioluminescence imaging. Surprisingly, LNP-GD2 mAb demonstrated reduced primary tumor uptake and preferential accumulation in the liver (Fig. 2C). Suspecting the diminished efficacy of LNP-GD2 mAb could be due, in part, to the increase in particle size or steric hinderance due to the high loading density of mAb on the LNP surface ^66^, we opted to replace full-length ch14.18 with a single-chain variable fragment (scFv) derivative ^67^ (Fig. 2D). We designed our scFv with a long flexible (GGGS)_4_ linker between the variable light (VL) and variable heavy (VH) chains, and a free C-terminal cysteine to facilitate efficient thiol-maleimide conjugation (Fig. 2D). Post-production validation of scFv activity was carried out by immunostaining and subsequent flow cytometric analysis demonstrating specificity comparable to ch14.18 (Fig. 2E). Successful conjugation was determined by SEC and DLS demonstrating a LNP diameter increase of ∼5 nm following scFv conjugation (Fig. 2F, Sup. Fig. 1A & B). Next, LNP-GD2 scFv conjugates carrying GLuc mRNA (3 mg/kg) were administered intravenously to mice harboring orthotopically engrafted neuroblastoma tumors. Organs were harvested 24-hours post-injection to assess GLuc expression, by means of BLI, in the liver, primary tumor, contralateral kidney, heart, lungs, and spleen by *ex vivo* bioluminescence imaging. Encouragingly, LNP-GD2 scFv demonstrated a narrowed biodistribution with appreciable primary tumor and liver uptake (Fig. 2G).

Offering greater resolution over bulk tissue analysis, we next assessed neuroblastoma cell-specific uptake, comparing unconjugated LNPs, LNP-GD2 mAb, and LNP-GD2 scFv. For these experiments, we encapsulated GFP mRNA. No appreciable change in particle size following encapsulation was observed when transitioning from GLuc to GFP mRNA (Fig. 3A). LNPs carrying GFP mRNA (3 mg/kg mRNA) were administered intravenously. Organs were harvested after a 24-hour incubation period and primary tumors were recovered. Single-cell suspensions were prepared from primary tumors and subsequently analyzed for GFP expression by flow cytometry (Fig. 3B & C). LNP-GD2 scFv demonstrated significantly higher neuroblastoma-specific uptake (47%, *P* < 0.002) than unconjugated LNP (30%) or LNP-GD2 mAb (28%; Fig. 3D). As such, we selected LNP-GD2 scFv as the initial carrier system for the systemic delivery of Cas9^D10A^ mRNA *in vivo*.

**Figure 3.**
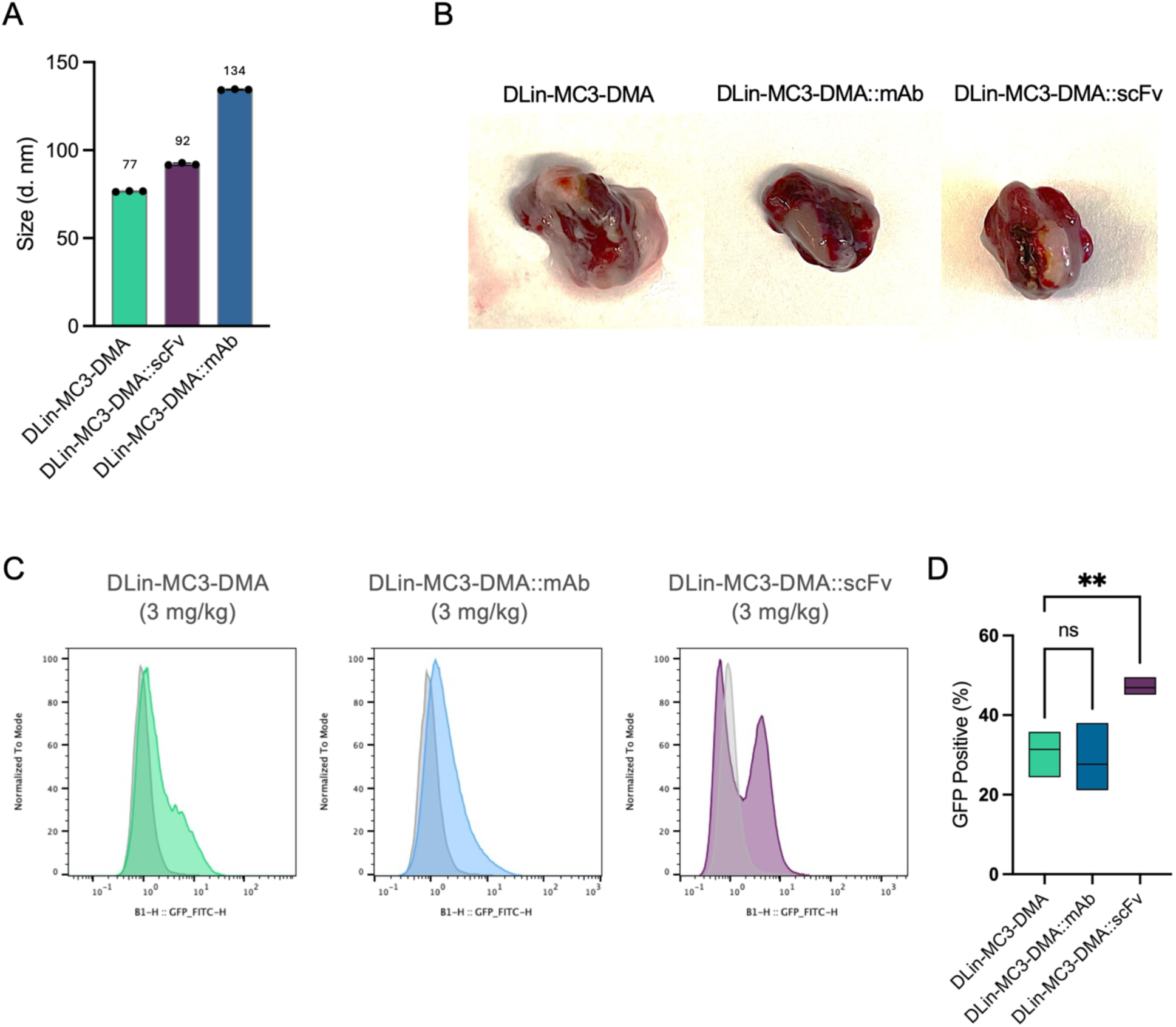
Functionalization of DLin-MC3-DMA LNPs with anti-GD2 scFv improves neuroblastoma-specific uptake. **A)** Average size (Z) of DLin-MC3-DMA, DLin-MC3-DMA – anti-GD2 antibody conjugated (DLin-MC3-DMA::mAb), and DLin-MC3-DMA – anti-GD2 scFv conjugated (DLin-MC3-DMA::scFv) LNPs encapsulating GFP mRNA (3 mg/kg) as determined by DLS. Data is presented as mean ± s.d. **B)** Representative images of neuroblastoma primary tumors used to analyze cell-specific expression of GFP. **C – D)** Flow cytometric analysis of GFP expression in dissociated primary tumor cells at 24-hours post-delivery of DLin-MC3-DMA (n = 3 mice), DLin-MC3-DMA::mAb (n = 5 mice), and DLin-MC3-DMA::scFv (n = 5 mice). Data are presented with a line indicating the mean. Data were analyzed using a one-way ANOVA with Dunnett’s multiple comparisons test; ns, P > 0.05; **, P ≤ 0.01.

### Cas9^D10A^ ameliorates disease burden in disseminated tumor model

For the analysis of Cas9 nickase efficacy *in vivo*, we continued to use SK-N-BE(2)C cells, as they have with an average *MYCN* copy number of 550 ^32^. These cells were modified to express, as stable transgenes, firefly luciferase to permit monitoring of tumor burden and one of two single guide RNA (sgRNA) to evaluate Cas9 nickase specificity: our therapeutic sgRNA (sgMYCN) ^32^, which targets a non-coding region downstream of *MYCN*, or our control sgRNA (sgAAVS1), which targets a non-amplified safe harbor locus corresponding to *PPP1R12C* – AAVS1 ^68^. In SK-N-BE(2)C cells, Cas9^D10A^ programmed with sgMYCN elicits substantial toxicity *in vitro*, whereas Cas9^D10A^ programmed with sgAAVS1 does not ^32^.

Given that oncogene amplification is associated with metastatic disease ^4,41–43,69^, we replaced our orthotopic neuroblastoma mouse model with a model of disseminated disease. As such, NSG mice were injected intravenously with SK-N-BE(2)C cells expressing either *MYCN* or *AAVS1* targeting sgRNA (Fig. 4A). Successful tumor engraftment was confirmed at 7-days post-injection by *in vivo* bioluminescence imaging, which demonstrated progressive disease in the liver and, in some instances, the bone marrow (Fig. 4B). Following confirmation of tumor engraftment, we initiated a 14-day treatment regimen. Here, LNP-GD2 scFv conjugates carrying Cas9^D10A^ mRNA (1.5 mg/kg mRNA) or a vehicle control were administered intravenously once every 24 hours for 3 consecutive days, and then once every 48 hours for the remainder of the 14-day treatment window for a total of 8 injections (Fig. 4A). LNP-GD2 scFv carrying Cas9^D10A^ mRNA demonstrated an average size (Z) of 79 nm and PDI of 0.169 (Fig. 4C & D). Disease progression was monitored once a week by *in vivo* bioluminescence imaging until the end of the treatment.

**Figure 4.**
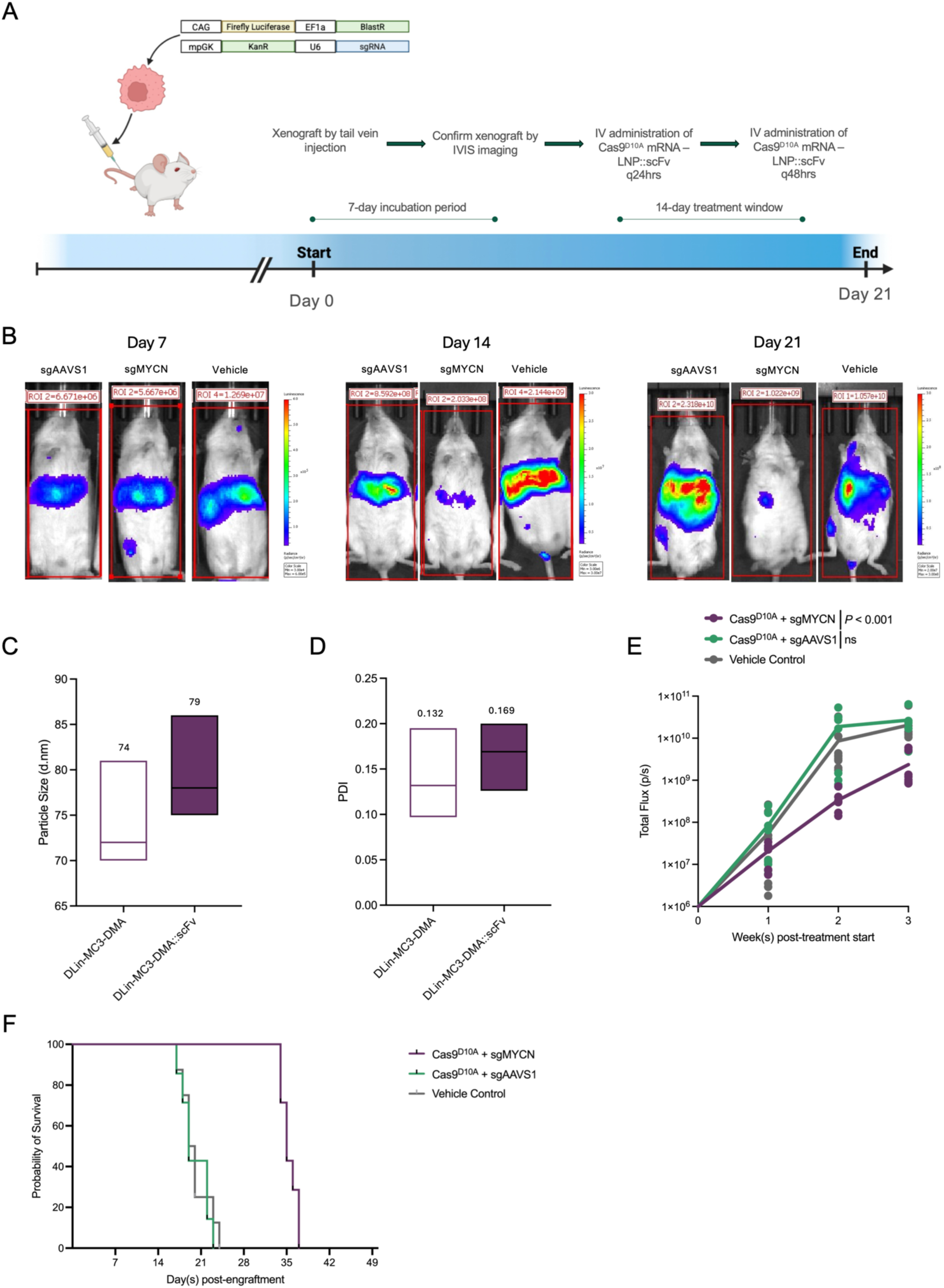
Cas9^D10A^ – mediated cell-killing demonstrates therapeutic effects *in vivo*. **A)** Experimental design to assess the therapeutic effects of Cas9^D10A^ – mediated cell-killing in a model of disseminated neuroblastoma. SK-N-BE(2)C cells harboring transgenes expressing Firefly luciferase and an sgRNA (sgMYCN or sgAAVS1) were injected systemically into NSG mice. After 7 days, LNP-GD2 scFv conjugates carrying Cas9^D10A^ mRNA (1.5 mg/kg mRNA) or a vehicle control are administered intravenously once every (q) 24 hours for 3 consecutive days, and then once every (q) 48 hours for the remainder of the 14-day treatment window for a total of 8 injections. **B)** Representative *in vivo* bioluminescence images of mice treated with Cas9^D10A^ targeting *MYCN* or *AAVS1*, or a vehicle control at 7-, 14-, and 21-days post-engraftment corresponding to 0-, 7-, and 14-days post-treatment initiation, respectively. **C – D)** Average size (Z-avg) and PDI of DLin-MC3-DMA and DLin-MC3-DMA – scFv conjugated (DLin-MC3-DMA::scFv) LNPs encapsulating Cas9^D10A^ mRNA (1.5 mg/kg) as determined by DLS (n = 7 independent replicates). Data are presented with a line indicating the mean. **E)** Bioluminescence intensities (BLI) of mice treated with Cas9^D10A^ targeting *MYCN* (n = 7 mice) or *AAVS1* (n = 7 mice), or a vehicle control (n = 11 mice). Changes in BLI assessed once a week during the treatment period. Data for each mouse are presented as individual data points. Significance was determined by two-way ANOVA with a Dunnett’s multiple comparison test relative to a vehicle control. **F)** Kaplan-Meier survival curve corresponding to mice in panel E.

At the treatment endpoint, Cas9^D10A^ targeting *MYCN* achieved a 9-fold reduction (*P* < 0.001) in total flux (p/s), whereas Cas9^D10A^ targeting *AAVS1* demonstrated no significant change in flux relative to a vehicle control (Fig. 4E). Additionally, the treatment regimen was well-tolerated with no appreciable fluctuations in body mass (Sup. Fig. 2) or adverse events observed. After the treatment regimen concluded, mice were monitored for overall survival until humane endpoint criteria were met. When targeting *MYCN* with Cas9^D10A^, mice demonstrated a 77% increase in mean survival, whereas targeting *AAVS1* resulted in no improvement in mean survival relative to a vehicle control (Fig. 4F).

### In vivo efficacy of Cas9^D10A^ can be augmented by CHK1 inhibition

CHK1 inhibitors have been developed to enhance the efficacy of conventional chemotherapeutics by interfering with DNA repair and cell-cycle control ^70–72^. Previously, we demonstrated that supplementation of Cas9^D10A^ with a subtherapeutic concentration of CHK1 inhibitor, MK8776 (SCH900776), significantly improves Cas9^D10A^ – mediated cell-killing in vitro ^32^. As such, we opted to investigate this combination therapy to augment the efficacy of Cas9^D10A^ – mediated cell-killing *in vivo*. Here, we modified the existing 14-day treatment plan with Cas9^D10A^ LNP-GD2 scFv to incorporate 3 cycles of MK8776, which was administered intraperitoneally (IP) at 7-, 14-, and 21-days post-engraftment at a subtherapeutic dose (10 mg/kg) ^73^.

Consistent with our observations *in vitro* ^32^, targeting *MYCN* with Cas9^D10A^ in combination with MK8776 achieved a 26-fold reduction (*P* < 0.0001) in total flux (p/s) at treatment end, whereas targeting *AAVS1* in combination with MK8776 or using MK8776 alone demonstrated no significant change in total flux relative to a vehicle control (Fig. 5A). Furthermore, when targeting *MYCN* with Cas9^D10A^ in combination with MK8776, mice demonstrated a 118% increase in mean survival relative to a vehicle control, whereas targeting *AAVS1* with Cas9^D10A^ in combination with MK8776 or administering MK8776 alone demonstrated no appreciable change in mean survival (Fig. 5B). No change in body mass or treatment tolerability was observed following the addition of MK8776 to the Cas9^D10A^ treatment regimen (Sup. Fig. 2).

**Figure 5.**
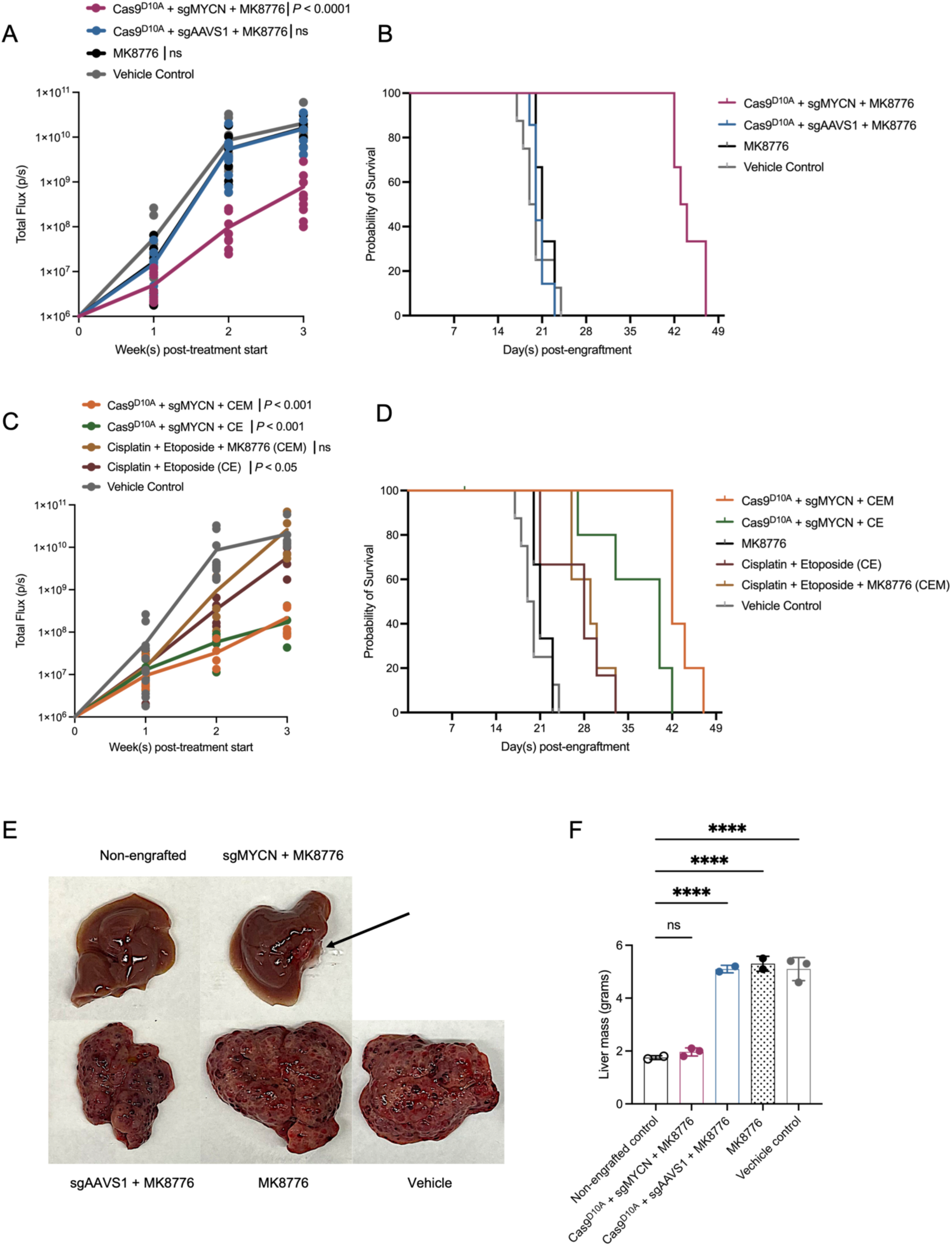
Cas9^D10A^ – mediated cell-killing can be augmented to increase therapeutic effects *in vivo*. **A)** Bioluminescence intensities (BLI) of mice treated with Cas9^D10A^ targeting *MYCN* (n = 9 mice) or *AAVS1* (n = 10 mice) and supplemented with MK8776, MK8776 only (n = 9 mice), or a vehicle control (n = 11 mice). Changes in BLI assessed once a week during the treatment period. Data for each mouse are presented as individual data points. Significance was determined by two-way ANOVA with a Dunnett’s multiple comparison test relative to a vehicle control. **B)** Kaplan-Meier survival curve corresponding to mice in panel A. **C)** Bioluminescence intensities (BLI) of mice treated with Cas9^D10A^ targeting *MYCN* and supplemented with cisplatin, etoposide, and MK8776 (CEM; n = 5 mice), Cas9^D10A^ targeting *MYCN* and supplemented with cisplatin and etoposide (CE; n = 6 mice), CEM only (n = 5 mice), CE only (n = 5 mice) or a vehicle control (n = 11 mice). Changes in BLI assessed once a week during the treatment period. Data for each mouse are presented as individual data points. Significance was determined by two-way ANOVA with a Dunnett’s multiple comparison test relative to a vehicle control. **D)** Kaplan-Meier survival curve corresponding to mice in panel C. **E)** Representative images of livers harvested from mice treated with Cas9^D10A^ targeting *MYCN* (n = 3 mice) or *AAVS1* (n = 3 mice) and supplemented with MK8776, MK8776 only (n = 3 mice), or a vehicle control (n = 3). Livers were harvested on the final day of treatment. Liver from non-engrafted control mice shown for reference. Disease mass in representative image of liver harvested from mice treated with Cas9^D10A^ targeting *MYCN* + MK8776 indicated with black arrow. **F)** Differences in liver mass corresponding to mice treated in panel E. Data are presented as individual data points around the mean ± s.d. Data were analyzed using a one-way ANOVA with Dunnett’s multiple comparisons test; ns, P > 0.05; ****, P ≤ 0.0001.

In addition to substantial *MYCN*-amplification, SK-N-BE(2)C cells harbor an inactivating missense mutation in *TP53* (C135F), rendering them less sensitive to genotoxic drugs ^74–76^. As such, we investigated if targeting *MYCN* with Cas9^D10A^ ± MK8776 could augment the efficacy of conventional chemotherapies. Here, we modified the existing 14-day treatment plan to incorporate a cocktail of cisplatin (3 mg/kg) and etoposide (30 mg/kg). The concentrations of etoposide and cisplatin administered are approximately half the maximum tolerated dose in mice ^77^. Cisplatin-etoposide ± MK8776 was administered intraperitoneally (IP) for a total of 3 cycles at 7-, 14-, and 21-days post-engraftment.

At treatment end, targeting *MYCN* with Cas9^D10A^ in combination with cisplatin-etoposide (CE) or cisplatin-etoposide and MK8776 (CEM) achieved 125-fold and 91-fold reductions (*P* < 0.001) in total flux (p/s), respectively, relative to a vehicle control (Fig. 5C). At the indicated concentrations, cisplatin-etoposide alone demonstrated 3.6-fold reduction (*P* < 0.05) in total flux (p/s), whereas cisplatin-etoposide and MK8776 demonstrated no significant reduction in total flux relative to a vehicle control (Fig. 5C). Furthermore, when targeting *MYCN* with Cas9^D10A^ in combination with cisplatin-etoposide, or cisplatin-etoposide-MK8776, mice achieved a 72% and 107% increase in mean survival relative to a vehicle control, respectively (Fig. 5D), whereas cisplatin-etoposide or cisplatin-etoposide-MK8776 alone achieved a 25% and 37% increase in survivorship, respectively, relative to a vehicle control (Fig. 5D). No significant fluctuation in body mass was observed (Sup. Fig. 2).

Targeting *MYCN* with Cas9^D10A^ in combination with MK8776 demonstrated the greatest improvement in mean survival relative to a vehicle control (Figure 6A & B). As such, we treated an additional cohort of mice harboring disseminated SK-N-BE(2)C tumors with Cas9^D10A^ targeting *MYCN* or *AAVS1* in combination with MK8776, MK8776 only, or a vehicle control and harvested their livers on the final day of treatment for a visual assessment of disease burden, *ex vivo*, relative to non-engrafted, healthy mice. In this experiment, targeting *MYCN* with Cas9^D10A^ in combination with MK8776 achieved substantial inhibition of disease progression in the liver (Fig. 5E). No significant increase in liver mass was observed in mice treated with Cas9^D10A^ targeting *MYCN* in combination with MK8776, whereas mice treated with Cas9^D10A^ targeting *AAVS1* in combination with MK8776, MK8776 only, or a vehicle control each demonstrated a ∼3-fold increase in liver mass, relative to a healthy mouse liver (Fig. 5F).

**Figure 6.**
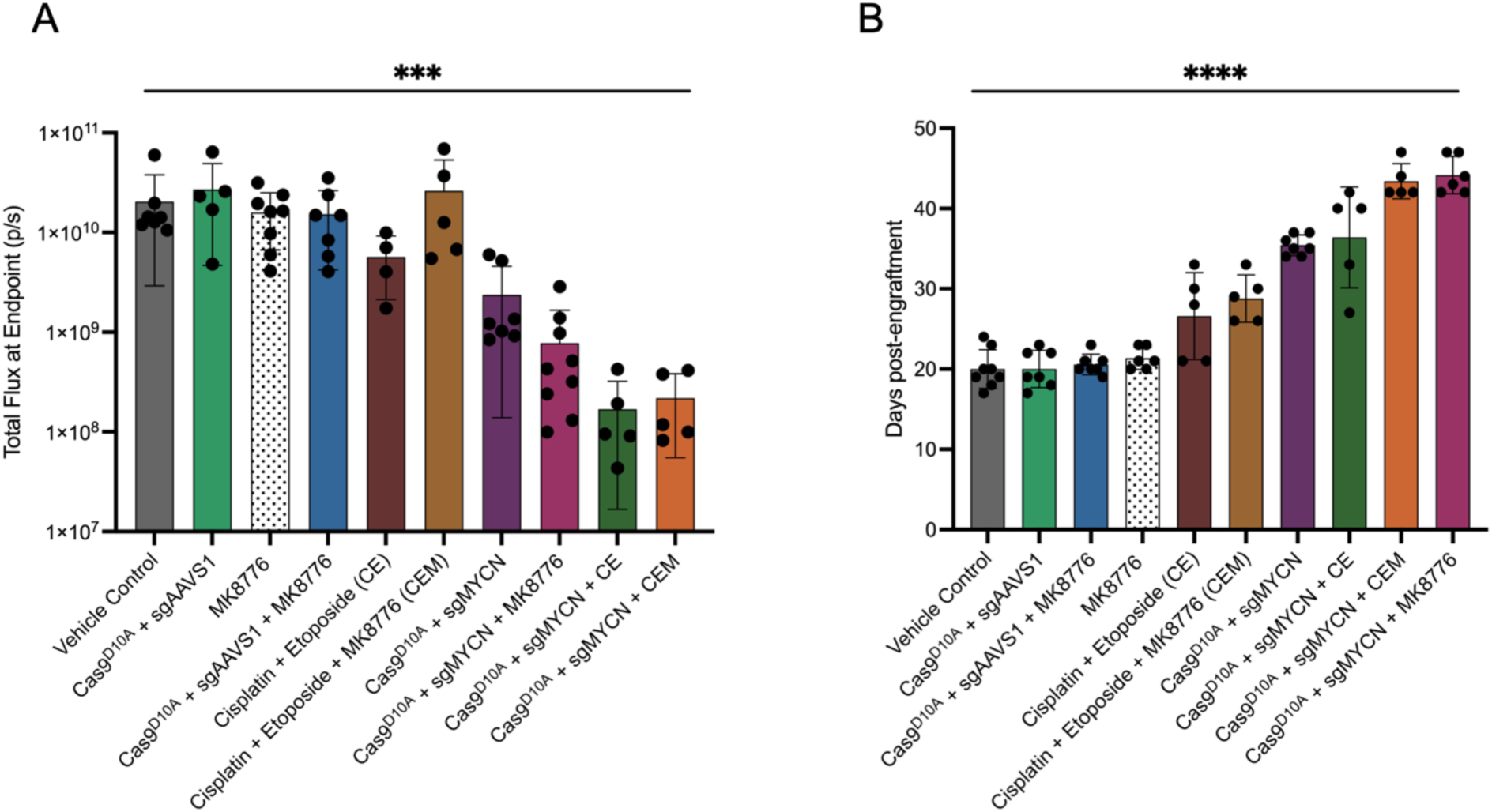
Cumulative comparison of change in BLI and survivorship for each treatment cohort. **A – B)** BLI at treatment end point (21-days post-engraftment; A) and survivorship (B) across all tested conditions. Data for each mouse are presented as individual data points around the mean ± s.d. Data analyzed using a one-way ANOVA; ***, P ≤ 0.001; ****, P ≤ 0.0001.

## DISCUSSION

Expanding on our previous work ^32^, we have demonstrated an initial framework for the development of a novel CRISPR–based therapeutic platform that selectively eliminates cancer cells by targeting gene amplifications. Gene amplification has been observed in 40 – 50% of cancers ^4,78^, but is notably frequent in breast and lung cancers, as well as neuroblastomas ^24,79,80^. Furthermore, gene amplification is often associated with relapsed and metastatic disease ^4,41–43^ for which novel therapeutics are desperately needed.

Our approach exploits gene amplifications as a critical, disease-specific vulnerability, while minimizing the potential for unwanted, non-specific genotoxicity in healthy tissues. Here, leveraging Cas9^D10A^ against gene amplifications generates large numbers of SSBs, which are subsequently converted into cytotoxic single-ended or double-ended double-strand breaks upon collision with the replisome during DNA replication ^32,33,36,39,40^. A significant advantage to this approach is its repeatability. Unlike Cas9 nuclease, Cas9 nickase produces few indels when targeted to a site within the genome ^81^. For example, targeting *MYCN* with Cas9^D10A^ in *MYCN*-amplified neuroblastoma cell lines resulted in indel rate of ≤ 3.7% within the surviving cell population ^32^. Low indel frequency minimizes the risk of resistance by virtue of target-sequence alteration, permitting repetitive dosing with the same sgRNA, which overcomes a potential resistance mechanism observed with Cas9 nuclease – based therapeutic approaches ^82–84^. Likewise, the use of Cas9 nickase instead of Cas9 nuclease minimizes the likelihood of damage to the genome of copy-normal cells should delivery occur to other tissues. Another significant advantage of this approach is its intrinsic modularity which offers a highly customizable platform suitable for personalized medicine. Lipid nanoparticles are versatile allowing for the modification of both passive and active targeting strategies, or the utilization of selective organ targeting molecules (SORT) ^49^ to enhance tissue-specific uptake. Likewise, the oncogenic landscape varies by cancer type, such that different oncogenes (*e.g*. *MYC*, *MYCN*, *ALK*, *ERBB2* (HER2), *EGFR*, and *CCND1*) may be preferentially amplified depending on the type of cancer. Importantly, our previous work indicates that Cas9^D10A^ – mediated cell-killing is strictly amplification dependent but independent of oncogene function, conferring efficacy in *ERBB2* (HER2) – amplified breast cancer and *MYC*-amplified lung and colorectal cancer *in vitro* ^32^. As such, Cas9^D10A^ can be programmed to target any amplified region by proper choice of the sgRNA. Finally, with growing interest in the development of combination therapies ^85^ with increased efficacy and reduced long-term, treatment-associated toxicities ^86,87^, we have demonstrated that the Cas9 nickase platform works cooperatively with other anti-cancer agents at less intensive or subtherapeutic doses to augment efficacy *in vivo* (Fig. 6A & B). Going forward, future efforts should place an emphasis on identifying optimal treatment dosage and scheduling to maximize efficacy.

Despite the encouraging observations made in this preliminary research there are limitations to this study. For ease of screening during this initial assessment of therapeutic efficacy, we employed transgenic cells modified for the stable, endogenous expression of each sgRNA. Given the favorable effects observed, subsequent assessments should incorporate the use of exogenously delivered sgRNA in conjunction with Cas9^D10A^ mRNA. Heavily chemically modified sgRNA is essential to reduce turnover and maintain potent *in vivo* activity in the context of LNP delivery, such as those containing extensive phosphodiester backbone chemical modifications ^88,89^. In addition, the initial assessment of this therapeutic platform *in vivo* was carried out in immunocompromised mice. As such, immune responses that may either increase or decrease the anti-cancer efficacy ^90^ of this platform are not reflected in these preliminary results.

## METHODS

### Ethics statement

Our study was conducted in compliance with the provisions set forth in the PHS/NIH Guide for the Care and Use of Laboratory Animals. All animal experiments were conducted by trained personnel in accordance with protocols approved by the Institutional Animal Care and Use Committee (IACUC) at the University of Massachusetts Chan Medical School (IPROTO202300000078 ;PROTO202000095). Criteria to ensure a humane endpoint included meeting one of following; ≥20% weight loss, dehydration and/or lethargy that does not improve within 24-hours, advanced disease with a bioluminescence intensity (BLI) ≥1 X 10^11^, or severe abdominal distension. Euthanasia was conducted with strict adherence to approved methods.

### Mammalian cell culture

SK-N-BE(2)C [CRL-2268] and HEK293T [CRL-3216] cell lines were sourced through the American Type Culture Collection (ATCC). All mammalian cell cultures were maintained at 37°C, 5% CO_2_, and ∼95% relative humidity. SK-N-BE(2)C cells were grown and maintained in Dulbecco’s Modified Eagle Medium/Nutrient Mixture F-12, GlutaMAX™ supplement (DMEM/F12, GlutaMAX™ supplement; Gibco, #10565018), supplemented with 10% fetal bovine serum (FBS; R&D Systems, #S12450H) and 1% penicillin-streptomycin (PS; 10,000 U/mL; Gibco, #15140122). HEK293T human embryonic kidney cells were grown and maintained in Dulbecco’s Modified Eagle Medium (DMEM; Gibco, #11965092) supplemented with 10% FBS (R&D Systems, #S12450H) and 1% PS (10,000 U/mL; Gibco, #15140122). Transgenic SK-N-BE(2)C cells were grown and maintained in the same media as previously described and further supplemented with Geneticin™ (1 mg/mL) (G418 Sulfate; 50 mg/mL; Gibco, #10131027) and/or blasticidin S HCl (10 µg/mL) (Gibco, #A1113903). Cell cultures were grown and maintained in 25cm^2^ (Celltreat, #229331) or 75cm^2^ (Celltreat, #229341) tissue culture flasks and tested for mycoplasma contamination at regular intervals using the e-Myco^™^ VALiD mycoplasma PCR detection kit (LiliF Diagnostics, #25239).

### Lipid nanoparticle formulation

The following components were purchased from MedChemExpress; DLin-MC3-DMA (HY-112251), DOTAP (HY-112754A), 306-O12B (HY-W590532), DMG-PEG 2000 (HY-112764), DOPC (HY-113424A), and DSPE-PEG-Maleimide (HY-140740). The following items were purchased from Millipore Sigma; DSPC (P1138) and cholesterol (C8667). Lipids were dissolved in 100% ethanol. Ionizable (DLin-MC3-DMA; 306-O12B) or cationic (DOTAP), helper (DSPC; DOPC), cholesterol, and PEGylated (DMG-PEG 2000) were combined at molar ratios of 50/38.5/10/1.5. When incorporating DSPE-PEG-Maleimide molar ratios were adjusted to 50/38.5/10/1.2/0.3.

Lipids were mixed with mRNA (50 mM citrate buffer, pH 4.5) at a 1:3 ratio using a NanoAssemblr Ignite (Precision NanoSystems Inc.) microfluidic system at a flow rate of 12 mL/min. LNPs were dialyzed against PBS (1X, pH 7.4) overnight at 4°C to remove residual ethanol. LNPs were concentrated using a Amicon Ultra Centrifugal Filter (100 kDa MWCO; MilliporeSigma) with subsequent buffer exchange into PBS (1X; 12% sucrose (w/v); pH 7.4) for storage at -80°C ^91^. Particle size and PDI were measured by DLS using a Zetasizer Nano (Malvern Panalytical). Encapsulation efficiency and mRNA concentration was measured by RiboGreen assay (ThermoFisher Scientific, R11491).

### In vitro transcription

*In vitro* transcription vectors incorporate a T7 RNA polymerase promoter for the production of synthetic GLuc, GFP, and Cas9^D10A^ mRNA. Synthetic mRNA products incorporate 5’ and 3’ UTRs derived from human hemoglobin alpha (hHBα) with a Kozak sequence, and amino-terminal enhancer of split (AES) and mitochondrially encoded 12S rRNA (mt-RNR1) motifs, respectively, as previously described ^32,92–94^, and a poly-A_150_ tail ^32^. The Cas9^D10A^ coding sequence was codon optimized for mammalian expression and minimized uridine content. Essential uridine bases were replaced with the modified nucleobase N1-methylpseudouridine (m1Ψ; TriLink, N-1081), as previously described ^32,95^. *In vitro* transcription of synthetic mRNAs was performed using the HiScribe T7 High Yield RNA Synthesis Kit (New England Biolabs, E2040) as previously described ^32^. Removal of dsRNA contaminants was performed as previously described ^96^.

### Cell line development

Donor vectors were designed to facilitate integration by *piggyBac*-mediated transposition ^97^. Expression of firefly luciferase and mCherry from a single donor vector was mediated by the incorporation of a synthetic CAG promoter and a self-cleavable P2A peptide linker between the gene coding sequences ^98^. The firefly-P2A-mCherry expression cassette also incorporated blasticidin resistance gene (BSD) under the control of a separate EF1α promoter to facilitate selection. Cloning of firefly-P2A-mCerry donor vector was achieved by Gibson Assembly (New England Biolabs, #E2611) in accordance with the manufacturer’s instructions. Separate donor vectors were constructed for the expression of single guide RNA under the control of U6 promoter and contained a neomycin-kanamycin resistance gene under the control of murine phosphoglycerate kinase (mPGK) promoter to facilitate selection. Oligonucleotides used to construct sgRNA inserts corresponding to sgMYCN (5’ – CAATGGAGACCCCATATGGG – 3’) or sgAAVS1 (5’ – GTCCCCTCCACCCCACAGTG – 3’) were ordered from GENEWIZ (Azenta Life Science, Inc., USA). Cloning of sgRNA donor vectors was achieved as previously described ^32^. Stable transfection of SK-N-BE(2)C cells for the production of firefly luciferase-P2A-mCherry and sgRNA expressing cells lines was achieved by piggyBac transposition as previously described ^32^.

### In vivo models

All models were generated in immunodeficient NSG (NOD.Cg-*Prkdc^scid^ Il2rg^tm1Wjl^*/SzJ) mice. Mice were obtained from as a gift from Michael Brehm (University of Massachusetts Chan Medical School) and subsequently bred in-house. Mice were housed in a controlled environment under standard conditions and kept on 12-hour light/dark cycles. Mice were maintained on a medicated diet (SCIDS MD’s - Gamma Irradiated, Bio-Serv, S0443). Mice between 8 – 16 weeks of age were used for experiments. Both male and female mice were used for each experiment with a ratio of ∼1:1.3. All mice were randomized based on BLI prior to experiments. Mice were assessed for changes in body mass once a week for the duration of the experiment.

SK-N-BE(2)C cells used for xenografts were cultured under standard conditions as previously described. Primary tumor models were established by orthotopically injecting 1 X 10^6^ cells resuspended in 100 µl of PBS (1X, pH 7.4) within the renal capsule on the mouse’s left side. Primary tumors were left to develop for 10 – 12 days and confirmed by bioluminescence imaging using an IVIS Spectrum CT (Perkin-Elmer). Disseminated tumor models were established by intravenously injecting (tail vein) 5 X 10^6^ cells resuspended in 200 µl of PBS (1X, pH 7.4) and supplemented with heparin sodium (0.4 IU/g; Fisher Scientific, AC411210010) to reduce the risk of embolism and promote extrapulmonary dissemination ^99^. Disseminated tumors were left to develop for 7-days and confirmed by bioluminescence imaging using an IVIS Spectrum CT (Perkin-Elmer). For bioluminescence imaging, mice received D-luciferin (IP; 150 mg/kg; GoldBio, CAS# 115144-35-9) prior to visualization.

### Delivery of LNPs in vivo

All administrations of LNPs and LNP vehicle controls were conducted by slow intravenous injection of lateral tail veins using a 28-gauge needle (Fisher Scientific, 14-826-79). Mice received GLuc or GFP mRNA – LNPs at a concentration of 3 mg/kg mRNA. Mice received Cas9^D10A^ mRNA – LNPs at a concentration of 1.5 mg/kg mRNA. Injection volumes were maintained at 150 µL and supplemented with low-dose heparin (0.08 IU/g) to reduce the risk of embolism ^100,101^ in mice receiving LNPs, as well as vehicle only controls to control for the intrinsic anti-cancer properties of heparin ^102,103^ in mice.

### Ex vivo imaging

All mice received orthotopic xenografts by renal capsule injection to generate primary tumors. Mouse treatment cohorts were randomized by BLI measured at day 10 – 12 post-engraftment. Mice received GLuc mRNA – LNPs (IV; 3 mg/kg mRNA) before being harvested at 24-hours post-injection. Mouse liver, primary tumor, contralateral kidney, heart, lungs, and spleen were collected and stored in PBS (1X; pH 7.4) on ice. Mouse organs were washed with PBS (1X; pH 7.4) and then submerged in a coelenterazine solution (100 µM; ThermoFisher Scientific, J66823.MA) for 30 minutes at room temperature. Mouse organs were rinsed briefly with PBS (1X; pH 7.4) and then immediately imaged using an IVIS Spectrum CT (Perkin-Elmer).

### Expression and purification of single-chain variable fragments

Anti-GD2 single-chain variable fragments (scFv) were derived from the GD2-specific antibody ch14.18 (Dinutuximab) using variable light (VL) and heavy (VH) regions described previously ^67^. The coding sequence was cloned into the pET-21a bacterial expression vector comprising the VL-VH chains joined by a (GGGS)_4_ linker and flanked by a N-terminal T7-tag and a C-terminal 6xHis-tag. Coding sequences were bacterial codon optimized for expression in SHuffle T7 competent *E. coli* (New England Biolabs, C3026J). Seed cultures of SHuffle T7 competent *E. coli* were established in 20 mL of 2xYT growth media (Fisher Scientific, DF0440-17) supplemented with carbenicillin (Fisher Scientific, 10-177-012) and incubated overnight at 30°C, 250 RPM. Next, seed cultures were pelleted at 2500 x g for 10 minutes, to discard old media, and resuspended in fresh media. Expression cultures (1L) were inoculated with 5 mL of seed culture to produce an initial OD_600_ of 0.02 – 0.05 and incubated at 30°C, 250 RPM to an OD_600_ of 0.2. After reaching an OD_600_ of 0.2, incubation temperature was dropped to 16°C and growth continued to an OD_600_ of 0.4 before being induced with IPTG (0.5 mM) for 16 hours. Next, expression cultures were pelleted 3500 x g for 20 minutes and resuspended in Tris buffer (50 mM Tris-HCl, 20 mM imidazole, 300 mM NaCl, pH 8.0) with Halt protease inhibitor (1:200; ThermoFisher Scientific, 78439). Resuspended pellets were lysed mechanically using a LM20 Microfluidizer Processor (Microfluidics). Clarified lysates were centrifuged ≥20,000 x g to pellet inclusion bodies. Inclusion bodies were resuspended by sonication in denaturing Tris buffer (50 mM Tris-HCl, 300 mM NaCl, 6M GdnHCl, 1 mM TCEP, pH 8.0) and purified by Ni-NTA chromatography (Qiagen) under denaturing conditions. Isolation of scFv was confirmed by SDS-PAGE and western blot analysis (Sup. Fig. 3 A&B). Isolated scFv was cleaned by anion exchange chromatography (Capto Q-resin, Cytiva) to remove residual nucleic acids and endotoxin. Removal of nucleic acid and endotoxin was confirmed by UV-Vis spectroscopy and Pro-Q™ Emerald 300 Lipopolysaccharide Gel Stain Kit (ThermoFisher Scientific, P20495; Sup. Fig. 3C), respectively. Purified, denatured scFv was refolded by stepwise removal of GdnHCl as previously described ^104^. Stepwise removal of GdnHCl and buffer exchange were performed using an ÄKTA flux s tangential flow filtration (TFF) system. Refolded scFvs were stored in PBS (1X, 1 mM TCEP, 10% glycerol) at -80°C.

### SDS-PAGE and western blot analysis

For SDS-PAGE, scFv samples were resolved using 10 – 20% tricine gels (Invitrogen, #EC6625BOX) in tricine SDS running buffer (1X; Invitrogen, #LC1675) at 130V constant voltage for 1.5 hours at room temperature. Resultant gels were stained with SYPRO orange (Invitrogen, S6650) in accordance with the manufacturer’s instructions. For western blot, unstained gels were transferred to PVDF membranes using the iBlot Dry Blotting System and blocked for 1 hour at room temperature in SuperBlock Blocking Buffer (0.05% Tween-20; Thermo Scientific, #37535). Blots were incubated with anti-T7 antibody (1:1000, Bethyl Laboratories, A190-117A) overnight at 4°C with mild agitation. Blots were washed for 5 minutes in TBST for a total of 3 washes then incubated for 1.5 hours at room temperature with anti-rabbit IgG, HRP-conjugated antibody (1:3000, Cell Signaling Technology, #7074). Resultant blots were incubated in ECL substrate (Thermo Scientific, #32106) for 1 – 2 minutes before visualization using ChemiDoc Touch Imaging System (BioRad).

### Flow cytometry

All mice received orthotopic xenografts to develop primary tumors and were randomized by BLI into cohorts for treatment as previously described. Mice received GFP mRNA – LNPs (IV; 3 mg/kg mRNA) before being harvested at 24-hours post-injection. Primary tumors were collected and stored in PBS (1X; pH 7.4) on ice before being minced into small pieces (≤ 2mm) with a razor blade. Minced tumor tissue was then resuspended in 5 mL of tumor digestion media containing 500 µL of collagenase/hyaluronidase blend (STEMCELL Technologies, 07912), 750 µL of DNase I solution (1 mg/mL), and 3.75 mL of DMEM/F12, GlutaMAX medium and incubated for 25 minutes at 37°C with mild agitation. Digested tumor tissue was passed through a 40 µm nylon mesh cell strainer and pelleted at 300 x g for 10 minutes. Cell pellets were resuspended gently in 10 mL of ammonium-chloride-potassium (ACK) lysing buffer (Gibco, A1049201) and incubated for 5 minutes at room temperature to remove red blood cells. Tumor cells were pelleted at 300 x g for 10 minutes, resuspended in PBS (1X; pH 7.4) then analyzed for GFP expression using the MACSQuant Analyzer 10 Flow Cytometer (Miltenyi Biotec) and FlowJo analysis software.

For the validation of scFv activity, SK-N-BE(2)C (GD2+) and HEK293T (GD2-) cells were prepared in single-cell suspensions in PBS (1X; pH 7.4) and incubated with either anti-GD2 scFv (1 µg) or anti-GD2 mAb (5 µg) per 1 X 10^6^ cells for 1 hour. Cells incubated with anti-GD2 scFv were then incubated with anti-T7 antibody (1:100, Bethyl Laboratories, A190-117A) then with goat anti-rabbit IgG (H+L) AF488 (1:250; Invitrogen, A-11008) for 40 minutes each. Cells incubated with anti-GD2 mAb (Dinutuximab) were then incubated with goat anti-human IgG (H+L) AF488 (1:250; Invitrogen, A-11013) for 40 minutes. All incubations were performed at 4°C with mild agitation. All cells were washed twice with PBS (1X; pH 7.4) supplemented with 1% FBS between antibody incubations. Control cells were incubated with their respective secondary antibody only. Post-staining, cells were resuspended in PBS (1X; pH 7.4) then analyzed using the MACSQuant Analyzer 10 Flow Cytometer (Miltenyi Biotec) and FlowJo analysis software.

### Production of LNP-mAb and LNP-scFv conjugates

LNPs were prepared with DSPE-PEG-Maleimide to facilitate maleimide-thiol conjugation. Full-length anti-GD2 IgG (ch14.18; Dinutuximab) were modified with SATA (N-succinimidyl S-acetylthioacetate) to introduce sulfhydryl groups within the fragment crystallizable region (Fc). Modification of ch14.18 was performed using a Pierce™ Sulfhydryl Addition Kit (ThermoFisher Scientific, 23460) in accordance with the manufacturer’s instructions. Anti-GD2 scFv was designed with a free, C-terminal cysteine residue and did not require any further modification prior to coupling. SATA-modified antibodies, or scFv, were mixed with maleimide functionalized LNPs at a molar ratio of 2:1 (DSPE-PEG-Mal to scFv) in PBS (1X; 1 mM TCEP, pH 7.4) and incubated overnight at 4°C with mild agitation. Unbound antibody or scFv was removed by gel filtration using Sepharose CL-4B size exclusion resin (MilliporeSigma, CL4B200). Resultant LNP-mAb or LNP-scFv conjugates were concentrated using a Amicon Ultra Centrifugal Filter (100 kDa MWCO; MilliporeSigma) with subsequent buffer exchange into PBS (1X; 12% sucrose (w/v); pH 7.4) for storage at -80°C ^91^. Particle size and PDI was measured by DLS using a Zetasizer Nano (Malvern Panalytical).

### Combination treatment in disseminated neuroblastoma model

The following items were purchased from MedChemExpress; SCH900776 (MK8776; HY-15532), etoposide (HY-13629), and cisplatin (HY-17394). Combination treatments consisted of Cas9^D10A^ mRNA LNPs supplemented with MK8776 (10 mg/kg), cisplatin (3 mg/kg) and etoposide (30 mg/kg), or MK8776 (10 mg/kg), cisplatin (3 mg/kg) and etoposide (30 mg/kg). Some assessments of therapeutic efficacy (e.g. IVIS imaging to estimate tumor burden) were conducted within a 14-day treatment window. LNP-GD2 scFv carrying Cas9^D10A^ mRNA (1.5 mg/kg mRNA) were administered to mice intravenously in the lateral tail vein once daily for 3 consecutive days, then once every 48-hours until day 14 post-treatment start. MK8776, cisplatin-etoposide, and MK8776-cisplatin-etoposide) were administered intraperitoneally for a total of 3 cycles on post-engraftment day 7, 14, and 21. Injection volumes were maintained at 150 µL (IV) and 200 µL (IP). All mice, where applicable, received route-specific administration of a vehicle control. For example, mice receiving MK8776 (IP; 10 mg/kg) also received intravenous administration of a vehicle control. Mice receiving cisplatin-etoposide often presented with fur loss and mild dehydration secondary to fatigue and diarrhea, which improved with subcutaneous infusions of saline between treatment cycles.

### Statistics and Reproducibility

No statistical method was used to predetermine sample size. Mice used in experiments were randomized into cohorts based on BLI. The Investigators were not blinded to allocation during experiments and outcome assessment. No data were excluded from the analyses. All statistical analyses were carried out using GraphPad Prism (10.2.3) software. Quantitative data, apart from where individual values are demonstrated, represent the mean ± standard deviation. Comparative statistical analyses were conducted using ordinary one-way ANOVA or two-way ANOVA with a Tukey’s or Dunnett’s multiple comparison test where applicable; ns, P > 0.05; *, P ≤ 0.05; ** P ≤ 0.01; ***, P ≤ 0.001; ****, P ≤ 0.0001.

### Data Availability

The data generated in this study are available upon request from the corresponding author.

## ACKNOWLEDGEMENTS

We thank Kim Wigglesworth and Pamela St. Louis for their management and care of our mouse colony. We also thank former and current members of the Wolfe laboratory, as well as colleagues within the Molecular, Cell and Cancer Biology Department for helpful suggestions and discussions. We thank Michael Brehm and his laboratory for providing the initial NSG mice needed for the studies. We are grateful to UMass Chan’s BRIDGE Innovation & Business Development Fund for supporting this research.

## AUTHOR CONTRIBUTIONS

M.B.H. and S.A.W. conceptualized the project and designed the experiments. M.B.H. conducted the experiments. M.B.H. and S.A.W. interpreted the data and wrote the manuscript.

