## Supplementary Figures for "Systemic delivery of CRISPR-Cas9 nickase suppresses oncogene amplified cancer progression"

A

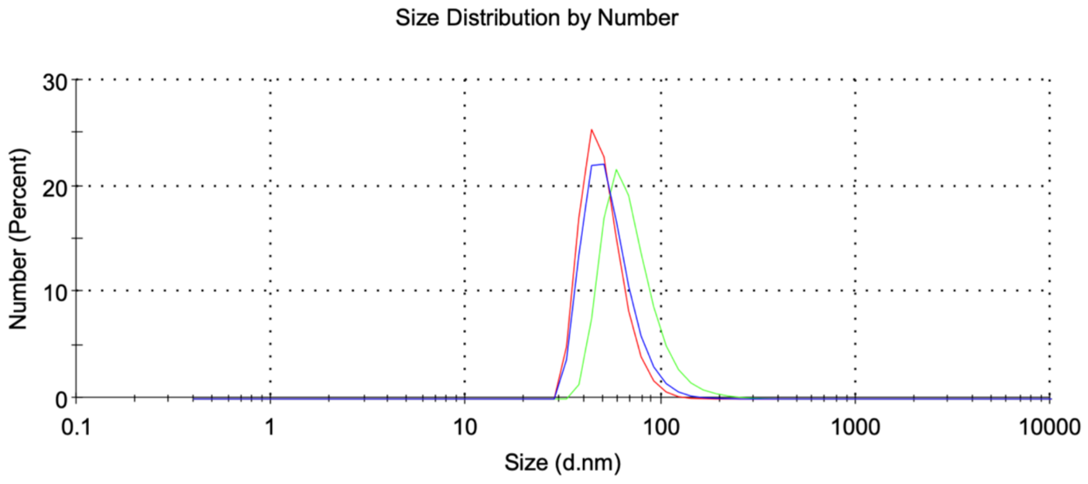


DLin-MC3-DMA; DLin-MC3-DMA::mAb; DLin-MC3-DMA::scFv

B

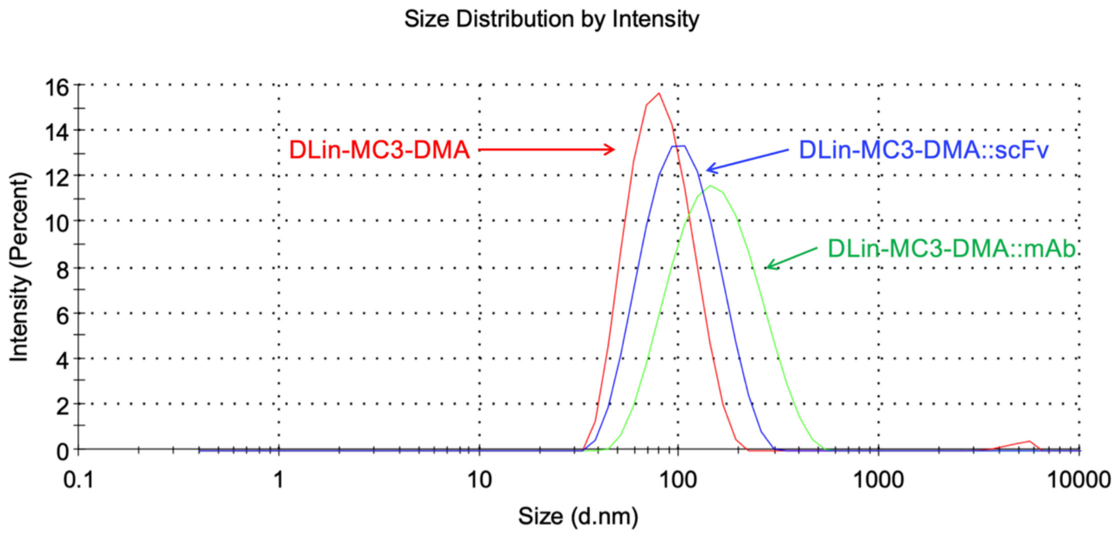

**Supplementary Figure 1. Confirmation of LNP – antibody or LNP – scFv conjugation post-purification by dynamic light scattering A)** Distribution of particle size by number that demonstrates the count of particles in different size bins. **B)** Distribution of particle size by intensity which demonstrates the amount of light scattered by particles of different sizes. **A – B**) Distributions represent increases in size comparable to known diameters of IgG and scFv post-purification.

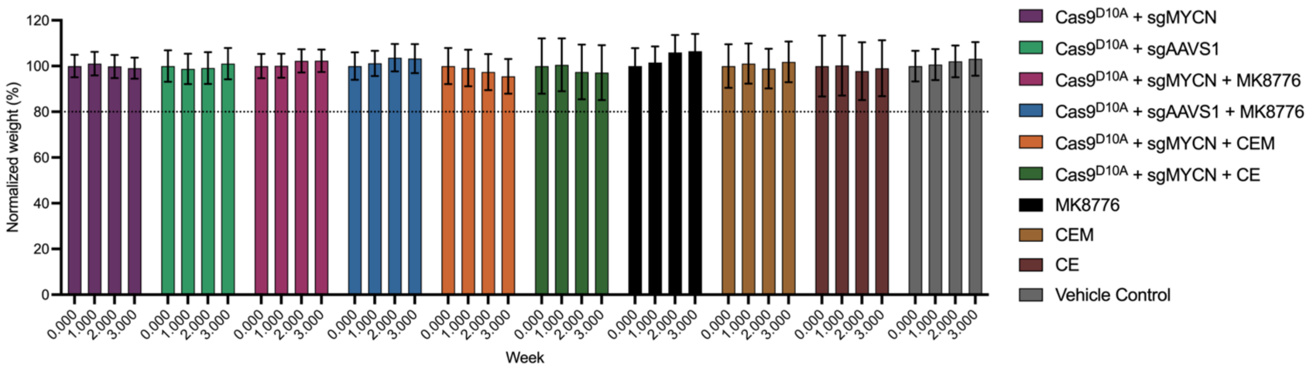

Normalized weight (%)
0
20
40
60
80
100
120
0.000
1.000
2.000
3.000
Week
Cas9D10A + sgMYCN
Cas9D10A + sgAAVS1
Cas9D10A + sgMYCN + MK8776
Cas9D10A + sgAAVS1 + MK8776
Cas9D10A + sgMYCN + CEM
Cas9D10A + sgMYCN + CE
MK8776
CEM
CE
Vehicle Control

**Supplementary Figure 2. Weekly body mass measurements for disseminated neuroblastoma mouse models under treatment.** All mice were weighed prior to engraftment and once a week thereafter for the duration of the treatment period. No mice were observed to lose ≥20% (black dotted line) of their body mass at any timepoint during the treatment period. Data has been normalized to display pre-treatment weights as 100% and are presented as mean ± s.e.m.

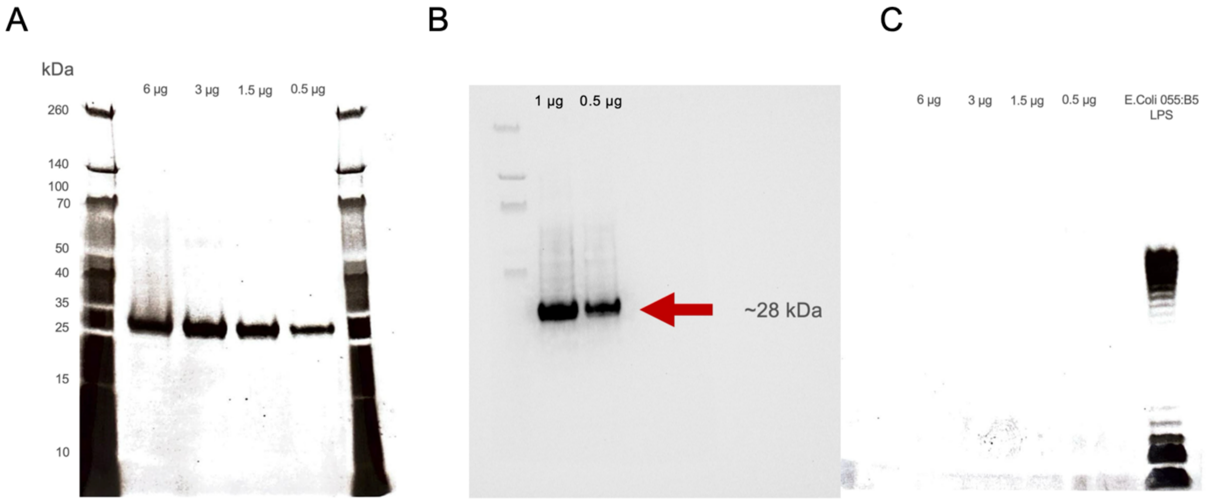
A
kDa
6 μg
3 μg
1.5 μg
0.5 μg
260
140
100
70
50
40
35
25
15
10
B
1 μg
0.5 μg
~28 kDa
C
6 μg
3 μg
1.5 μg
0.5 μg
E.Coli 055:B5
LPS

**Supplementary Figure 3. Isolation of anti-GD2 scFv for conjugation with LNPs. A)** SYPRO orange stained SDS-PAGE gel of anti-GD2 scFv (28 kDa) produced by prokaryotic expression in SHuffle T7 competent *E. coli*. SDS-PAGE analysis suggests a homogenous product with negligible contamination observed at increased concentrations. **B)** Western blot of anti-GD2 scFv produced by prokaryotic expression probing for N-terminal T7-tag. **C)** Post-purification assessment of residual endotoxin in anti-GD2 scFv aliquots prior to conjugation with LNPs. No endotoxin was detected at increasing concentrations of anti-GD2 scFv relative to a lipopolysaccharide (LPS) standard from Escherichia coli serotype 055:B5 (2.5 μg).
